# Cell-type specific analysis of heterogeneous methylation signal using a Bayesian model-based approach

**DOI:** 10.1101/682070

**Authors:** Daniel W Kennedy, Nicole M White, Miles C Benton, Rodney A Lea, Kerrie Mengersen

## Abstract

**Motivation:** Epigenome-wide studies are often performed using heterogeneous methylation samples, especially when there is no prior information as to which cell-types are disease associated. While much work has been done in ascertaining cell-type fractions and removing cell-type heterogeneity variation, relatively little work has been done in identifying cell-type specific variation in heterogeneous samples.

**Results:** In this paper, we present a Bayesian model-based approach for making cell-type specific inferences in heterogeneous settings, by using a logit-Normal sampling distribution and incorporating *a priori* knowledge of cell-type lineage. The method is applied to the detection of cell-type specific sex effects in methylation, where cell-type information is present as an independent verification of the results. Panels derived from this method contained more loci where CD8+T, CD19+B and Natural Killer cell-types were differentially methylated. The analysis suggests that an ensemble approach with this method included could be used for discovering cell-type specific methylation changes.

**Availability:** https://github.com/danwkenn/Bayes_CDM

## Introduction

Epigenome-wide Association Studies (EWASs) have been used extensively to find genomic loci where epigenetic changes are associated with some phenotype of interest. Epigenetic changes are commonly identified by differences in the DNA methylation levels of cells in accessible tissues such as blood. DNA methylation is a biological process whereby a methyl group (CH_3_) attaches to a Cytosine base which is proceeded by a Guanine base, referred to as a CpG locus. Methylation is known to vary between cell-types (1), and to play a role in cell differentiation (2) and normal cell function (3). Reinius et al. (1) were able to recover the haematopoietic lineage of whole blood immune cell-types using the methylation profiles of cell-sorted samples, indicating there are distinct cell-type and cell-type grouping methylation profiles.

The methylome has been found to have numerous associations with normal biological variation attributable to age (4) and sex (5, 6), as well as autoimmune disorders including multiple sclerosis (see Webb and de Arellano (7), Zulet et al. for recent reviews), diabetes (9), rheumatoid arthritis (10), and many types of cancer (11, 12). Additionally, there are known to be both phenotype- and disease-associated loci which only show association for specific cell-types. White et al. (13) used a Bayesian model selection approach to identify sets of panels with sex associations associated with celltypes as a function of cell lineage. It has been demonstrated that age has a measurably different effect on methylation for different cell-subtypes and tissue (4, 14), and Multiple Sclerosis is known to be specifically associated with T-cell differential methylation (15).

Many EWASs are conducted using samples from mixed celltype tissues, such as Whole Blood or PBMC samples, because of (a) the prohibitive cost of obtaining cell-sorted samples, (b) the lack of *a priori* knowledge for which cell-types are associated with the phenotype, and (c) the proliferation of heterogeneous data in the public domain. Given mixed cell samples are comprised of multiple constituent cell-types, mixed cell methylation data exhibit a profile which is a convolution of the profiles from the constituent cell-types. This gives rise to several issues when conducting an EWAS on mixed cell tissue samples. Firstly, differences in methylation between constituent cell-types introduce a large amount of variation which is unrelated to the phenotype of interest (16). Secondly, phenotype-related changes to the cell-type composition are manifested as methylation changes in mixed cell profiles at cell-type associated loci, and could lead to false associations at these loci. Thirdly, phenotype associations with less prevalent cell-types are less likely to be detected compared to that of more common cell-types.

Methylation data are often in the form of beta-values, each of which can be viewed as a measure of the probability that a strand of DNA in the sample is methylated at a given locus. Beta-values from mixed cell samples have been modelled as a linear combination of the underlying cell-type specific methylation levels, weighted by the cell-type proportions of the sample (17–19), an assumption referred to here as the linear mixing process, a term used to describe the equivalent process for gene expression data (20). While many algorithms have been developed which account for and remove variation associated with cell-type heterogeneity (21–23), there has been comparatively little research into finding and estimating effects of a phenotype or disease state on constituent cell-type methylation levels. This paper presents a statistical model for detecting and estimating differential methylation on the cell-type level in mixed cell samples.

One method of predicting associations at the cell-type level uses *a priori* known cell-type-specific regulatory information to suggest which cell-types are likely to be differentially methylated, given a set of phenotypically associated loci (24). However, since the method does not make this determination based on the data itself, it can only predict differential methylation for a given cell-type at loci that are previously known to be associated with that cell-type via regulatory information. Therefore, there is merit in a method which uses only the mixed-cell-derived data, with the greater potential of inferring the direction and size of the phenotype effects in each cell-type.

In this paper we propose a novel statistical model, which combined with an optimisation fitting procedure we refer to as Bayesian Cell-type level Differential Methylation (Bayes-CDM). The model preserves the linear mixing process assumption, but also implicitly enforces the boundary restrictions on the data and the parameters using a logit link function. Furthermore, this model incorporates prior information concerning relatedness of cell-type methylation profiles, based on the haematopoeitic lineage. It is known that methylation plays a role in cell-type differentiation, so the parameters governing cell-type methylation can be made to reflect this lineage via contrasts. None of the previous cell-type inference methods leverage this information, but in our method we establish a prior covariance structure to incorporate it. We therefore propose an extension of EWAS to the cell-type level, when only mixed cell samples are available.

The goal of an EWAS is to identify loci where there is an association between the methylation data and an underlying phenotype. The focus of this paper is to identify an association between the methylation level of a given cell-type’s methylation level and an underlying phenotype when only mixed cell data are available. Further, we assume that estimates of the cell-type proportions in each sample are available.

A simulation of mixed cell data is conducted to demonstrate the utility of the method in detecting cell-type level differential methylation. The method is then used for a combined data set of methylation samples from male and female subjects, with sex used as the phenotype to which related differential methylation is identified. Given that corresponding cell-sorted data was available, this provided a ground-truth for comparison with the output of Bayes-CDM and several competing methods.

## Methods

Let *I* be the number of methylation samples. Methylation data for a locus can be represented of an *I*-length vector **y** = (*y*_1_, …, *y*_*I*_)^T^ where *y*_*i*_ corresponds to the beta-value of the *i*^th^ sample. Beta-values are constrained to the unit interval, that is 0 ≤ *y*_*i*_ ≤ 1 for *i* = 1, …, *I*.

In this paper we consider a binary phenotype, whereby the *i*^th^ is assigned one of two possible phenotype levels, *δ*_*i*_ = 0, or *δ*_*i*_ = 1. The goal is to infer whether there is a difference in the methylation level between the two phenotype levels (0 and 1) for each constituent cell-type.

Given that the data are restricted to the unit interval, *y*_*i*_ is assumed to follow a logit-Normal distribution,

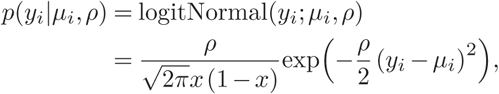

which is defined by the logit-median parameter *μ*_*i*_ ∈ ℝ and the precision parameter *ρ* ∈ ℝ_+_. The model is applied to each locus individually, meaning that for *I* samples, the likelihood is

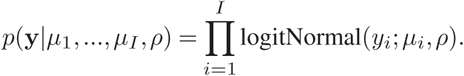

Several previous methods (18, 21, 25) used a Normal distribution for the beta-value. The mean value was specified as the linear combination of the underlying cell-type methylation values, weighted by their respective cell-type proportions. Since the mean and mode of the logit-Normal distribution are not available in closed form, the median of *y*_*i*_ was used.

Let *π*_*ik*_ be the cell-type proportion estimate for sample *i* and cell-type *k*, where *i* = 1, …, *I* and *k* = 1, …, *K*:

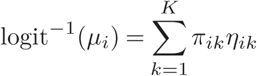

The median of *y*_*i*_ is therefore assumed to be the linear combination of the underlying cell-type methylation levels *η*_*i*1_, …, *η*_*iK*_, where each *η*_*ik*_ is constrained between 0 and 1. The cell-type proportion values are constrained to be positive, and values from each sample add to 1.

The linear predictor for each *η*_*ik*_ is parametrised in terms of a baseline *θ*_*k*_ and a shift *ϕ*_*k*_ for each cell-type.

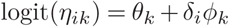

Therefore, *η*_*ik*_ can be thought of as the cell-type *k* methylation level in sample *i*. This parametrisation allows for the effect of phenotype on cell-type *k* methylation level to be estimated from *ϕ*_*k*_. When *ϕ*_*k*_ = 0, there is no difference between the methylation levels of cell-type *k* for the two phenotype levels. By having *θ*_*k*_ + *δ*_*i*_*ϕ*_*k*_ equal the logit of the cell-type methylation level, *θ*_*k*_ and *ϕ*_*k*_ do not need to be constrained in order for *η*_*k*_ and thus *μ*_*i*_ to remain on the unit interval.

We adopted a Bayesian modelling approach to overcome the issue of constrained data and parameters, and to incorporate cell-type lineage relationships as prior information. The model has 2*K* free parameters specifying the mean methylation level, but typical methylation data sets tend to have small sample sizes. Therefore, regularisation in the form of an informative prior distribution was used for the *θ* and *ϕ* parameters. A set of Normal shrinkage priors was placed on special linear combinations which incorporate the haematopoeitic lineage (see Figure 1). For the remainder of the paper we focus on the example of whole blood methylation data, which is composed of 6 major cell-types; CD14^+^ Monocyte, CD56^+^ Natural Killer (henseforth abbreviated to Monocyte and Natural Killer respectively), Neutrophil, CD19^+^B, CD4^+^ and CD8^+^T, however this method can be extended to any heterogeneous tissue.

**Fig. 1.**
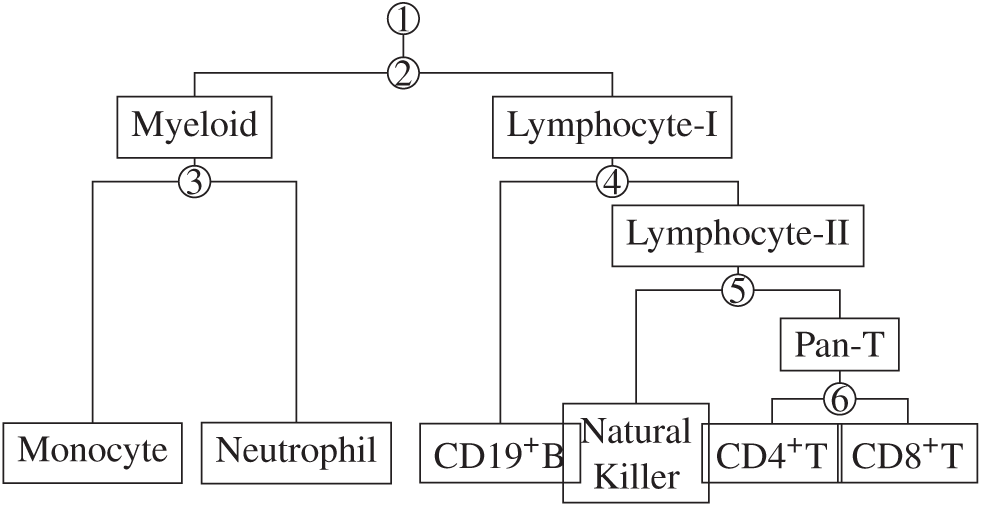
Cell-types and cell-type groupings as clustered by the haematopoietic lineage, with *nodes* each given a number. The nodes relate to the columns of the lineage matrix **A** while each of the cell-types relate to a row.

By associating a parameter with the node of the lineage rather than the cell-type, we can represent cell-type methylation as successive contrasts between the two cell-type groupings distinguished after each node. The phenotype level-0 cell-type methylation levels are parametrised by ***θ*** = (*θ*_1_, …, *θ*_*K*_)^T^. Let ***ξ*** = (*ξ*_1_, …, *ξ*_*K*_)^T^ be a set of contrast parameters associated with each node, such that ***ξ*** and ***θ*** are related via a lineage matrix **A**:

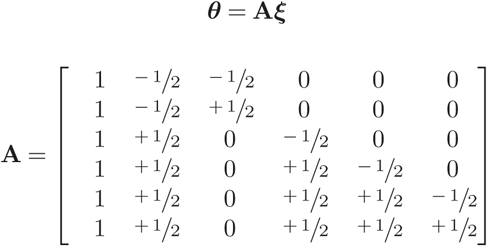

Each column relates to a node in the cell-type lineage, and each row refers to a cell-type. The corresponding mapping of contrast parameters to cell-type level parameters is given in Table 1. The parameters *ξ*_*q*_ differentiate between the cell-type groupings formed by the associated node *q*. For example, *ξ*_2_ acts as the contrast between Myeloid and Lymphocyte-I types, as each Myeloid cell-type methylation has a 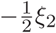 term, and each Lymphocyte-I cell-type baseline methylation has a 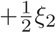 term. The subsequent *ξ* terms describe later cell-type differentiations in the lineage such as *ξ*_6_, which is the difference in baseline methylation between CD8^+^T and CD4^+^T. The exception is the first parameter *ξ*_1_ acts as an intercept, since it appears as a component in all constituent cell-types, and does not differentiate between cell-type groupings.

**Table 1.**
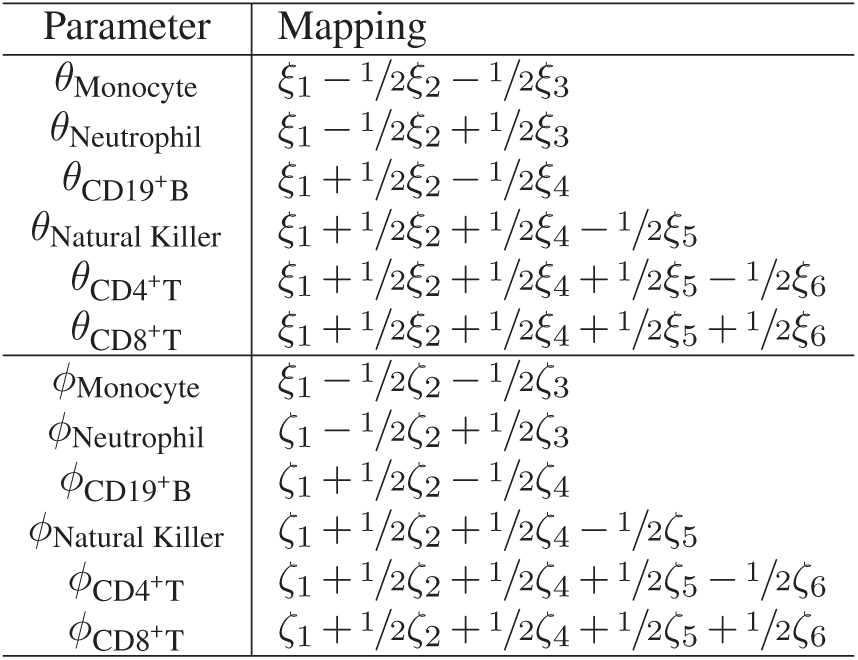
Mapping of contrast parameters *ξ* and *ζ* terms to the cell-type level parameters *θ* and *ϕ*.

The influence of cell-type lineage on the phenotype effect is also considered. In the same way as above, ***ϕ*** = (*ϕ*_1_, …, *ϕ*_*K*_)^T^ can be described in terms of node contrasts ***ζ*** = (*ζ*_1_, …, *ζ*_*K*_)^T^.

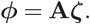

Here *ζ*_1_ acts as an overall shift in methylation level from phenotype level 0 to 1. The second parameter *ζ*_2_ is the difference in phenotype effect between Myeloid and Lymphocyte-I types, and so on.

We placed Normal priors with mean 0 on the linear contrast parameters, which have the effect of shrinking the posterior distributions and parameter estimates towards 0, and shrinking associated cell-type methylation level parameters (*θ* and *ϕ*) of related cell-types together. The degree of shrinkage is dependent on the precision of the prior distributions relative to the informativeness of the data as expressed by the likelihood. Therefore, as sample size and thus information from data increases, bias incurred from shrinkage decreases. Thus for *q* ∈ {2, …, *K*},

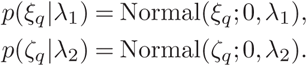

The prior distributions for the two precision parameters *λ*_1_ and *λ*_2_ were Gamma distributed with a shape parameter *α* = 1 and the shrinkage parameter *λ*_0_,

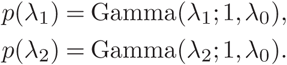

The mean of *p*(*λ*_1_) and *p*(*λ*_2_) is 1*/λ*_0_, so the effect of increasing *λ*_0_ is to decrease the prior mean of *λ*_1_ and *λ*_2_ and thereby increase the amount of regularisation. The value *λ*_0_ = 1 was chosen as a reasonable level of regularisation, given the high noise-low data context of methylation studies. It is straightforward to show that the marginal prior distribution of the *ξ* and *ζ* parameters is a standard Student’s *t* distribution with 2 degrees of freedom. For a simple empirical justification and investigation of the prior distributions of *θ* and *ϕ* parameters as a consequence of the above prior specification of *ξ* and *ζ* refer the Supplementary Material.

A half-Cauchy prior was used for the precision parameter *ρ*:

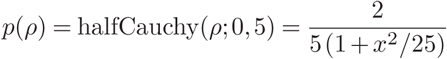

Since *ξ*_1_ and *ζ*_1_ are not cell-type related, we chose not to shrink them and instead use a weakly informative Cauchy distribution:

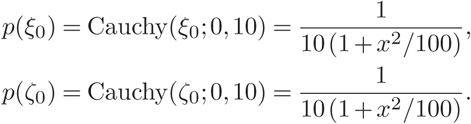

### A. Model Inference

The posterior densities for the model parameters were approximated using Laplace approximations. Maximum A Posteriori (MAP) estimates for the parameters 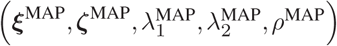 and an estimate of the Hessian matrix were calculated using numerical optimisation implemented in the STAN (26) software R package. To calculate posterior standard deviations of *ϕ* and *θ*, the standard deviation of the marginal distributions needed to be calculated from the corresponding Hessian estimate.

Since the parameter *ϕ*_*k*_ determined the effect size of phenotype on the methylation level of cell-type *k*, it is the basis for predicting if a given cell-type *k* is differentially methylated. Since we used a Laplacian approximation to the posterior, the probability

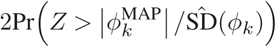

is a measure of the magnitude of the standardised effect size, where *Z* ∼ Normal(0, 1) and 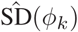 is the estimate of the standard deviation of the posterior for *ϕ*_*k*_. Consequently, we used this value with cut-off value *α* as a decision boundary. If this posterior probability for *ϕ*_*k*_ was less than *α*, then we predict that cell-type *k* is differentially methylated.

### B. Simulation study

We conducted a simulation study of mixed cell data to investigate the ability of the method to detect differential methylation in cell-types, in comparison to a number of other potential methods. An underlying set of cell-type methylation levels was constructed for four different scenarios (see Suppl. Figure 1) to investigate behaviour of detection methods over different profiles of differential methylation. We presumed a reasonable number of samples for a methylation study is *I* = 50, and precision parameter *ρ* = 30.

The four simulation scenarios were designed to investigate how varying the number of differentially methylated cell-types and the effect sizes affected the probability of detection (see Suppl. Figure 1 for visual depiction). Scenario 1 simulates a locus where the differential methylation is restricted to cell-types of the myeloid lineage grouping (Monocyte and Neutrophil), playing to the strength of Bayes-CDM, which uses the prior covariance structure to shrink related cell-type methylation levels together. Scenario 2 showed a similar situation, except the effect is much smaller. Scenario 3 showed a situation where differential methylation is exhibited in three cell-types, not all related. Finally, Scenario 4 showed a situation where only a single cell-type, the Monocyte, is differentially methylated. The method and a set of alternatives, described in Table 2, were used to predict if each cell-type was differentially methylated in all four scenarios. This was repeated for 100 simulations of each scenario to investigate the methods in terms of specificity and sensitivity.

**Table 2.** Alternative methods for finding differential methylation in cell-type heterogeneous samples. These methods represent four possible alternative approaches to making inferences; Orthogonal Effects Linear Regression (OE-LR); where only considering orthogonal effects to the cell-type proportion estimates; Aggregated Linear regression (Agg-LR), where each cell-type’s differential methylation is inferred separately; Popoulation-specific Expression Analysis (PSEA), which uses the full linear model and a subset selection method; and LASSO, which also uses the full model but uses the sparsity property of the regularising function in contrast with the PSEA method.

| Method | Abbreviation | Description |
| --- | --- | --- |
| Orthogonal Effects Linear Regression (standard heterogeneity-corrected linear regression) | OE-LR | Linear regression with cell-type proportion estimates included to account for cell-type heterogeneity, and a single orthogonal effect of phenotype. Model: $y_i = \delta_i \beta + \sum_{k=1}^K \pi_{ik} \gamma_k + \epsilon_i$ |
| Aggregated Linear Regression (extension of (18)) | Agg-LR | Method is run once for each cell-type. For cell-type $k$, the proportions of the other cell-types are aggregated together, and the following model is fitted: $y_i = \beta_0 + \beta_1 \pi_{ik} + \beta_2 \delta_i \pi_{ik} + \beta_3 \delta_i (1 - \pi_{ik}) + \epsilon_i$<br>The value of $\beta_2$ determines if cell-type differential methylation occurs. |
| Population-Specific Expression Analysis (25) | PSEA | Method uses a complete model with a phenotype effect for each cell-type: $y_i = \beta_0 + \sum_{k=1}^K \pi_{ik} \beta_k + \sum_{k=1}^K \pi_{ik} \delta_i \gamma_k + \epsilon_i$<br>The method then uses best subset selection based on the Akaike Information Criterion to select a subset of the coefficients. We set the cut-off value as 1 if the effect was not selected, and the standard $F$-test $p$-value from the OLS fit of the model if selected. |
| Least Absolute Shrinkage and Selection Operator (28) | LASSO | Method is a complete model as with PSEA, but applies a regularising penalty function on the coefficients which acts to select variables by shrinking some to 0. We only applied the shrinkage on the phenotype effect coefficients, and obtained a cut-off by choosing the value of the shrinkage parameter where the coefficient was shrunk to 0. |

In all scenarios, there was a non-phenotypic difference in the methylation levels between the Myeloid and Lymphocyte, as well as a difference between CD19^+^B and the other Lymphocyte types. In terms of the model, this translates to differing *θ*-values between cell-types. The Fluorescence-Activated Cell-Sorting (FACS) composition data from a mixed-sex data set (GSE88824) was used to fit Dirichlet parameters (via `diri.est` from the Compositional R package), and simulated composition data were drawn from this Dirichlet distribution.

Several alternative methods were used as a comparison for Bayes-CDM and are described in table 2. The Orthogonal Effects Linear Regression (OE-LR) method presented here models the phenotype effect as non-specific to cell-type, and as such should have a performance similar to methodologies which correct for cell-type heterogeneity without allowing for cell-types to have different phenotype effects. Some examples of this type of methodologies include (CITE,CITE, CITE), which all use the proportion estimation method by (27) to obtain proportion estimates. The Aggregated Linear Regression (Agg-LR) method represents a suggested extension of the method used by (18) for more than two cell-types. Here a separate model is fitted for each cell-type by aggregating other cell-type proportions together. Population-Specific Expression Analysis (PSEA) (25) and an implementation of the LASSO (28) represent the same, fully specified linear model, with a separate effect for each cell-type. They differ in that PSEA uses best-subset selection while LASSO uses *l*_1_-regularisation to select the optimal parameter set. In the case of the LASSO, the variable selection was limited to the interaction effects between cell-type fraction and phenotype, and 10-fold cross-validation was used to find the optimal tuning parameter for the LASSO. To the best of our knowledge, the LASSO has not been applied for this specific problem in methylation data, however given the interest in regularisation techniques we wanted to investigate how such a method would perform.

Since there are multiple cell-types; and therefore multiple possible cell-types with differential methylation, an ideal method correctly detects all cell-types with differential methylation. Consequently, the detection rate of differential methylation in each cell-type was calculated, as well as the rate of choosing five multi-cell-type measures described in Table 3. These measures were designed to quantify the simultaneous predictive performance over multiple cell-types.

**Table 3.** Measures for the evaluating predictive performance over multiple cell-types.

| Measure | Description |
| --- | --- |
| Predicted DM | Differential methylation detected in at-least one cell-type. |
| At least one true | Differential methylation correctly predicted in at least one cell-type. |
| At least one true, no false | Differential methylation correctly detected in at least one cell-type, without any incorrect detections. |
| All true associations | Differential methylation correctly detected for all cell-types with differential methylation. |
| Correct Subset | Differential methylation correctly detected for all cell-types with differential methylation, with no incorrect detections. |

### C. Case study: Finding Cell-type differential methylation associated with sex

Two publicly available data sets (GEO Accession Numbers: GSE35069 and GSE88824) containing both mixed cell and cell-sorted data, as well as Fluorescence Activated Cell Sorting (FACS) estimates of the cell-type proportions (see Supplementary Material of Reinius et al. (1)), were merged into a single combined data set. In total there were 5 female and 9 male subjects, which allowed us to identify differential methylation associated with sex using both the mixed cell data and independently using the cellsorted data.

#### C.1. Obtaining a ground-truth differential methylation using cell-sorted data

In this case study, each of the methods described in the previous section were used to predict which loci were differentially methylated for each cell-type from mixed-cell data. The cell-sorted data available provided an independent and accurate means of detecting differential methylation, and in consequence the differential methylation status detected via cell-sorted data were used as the groundtruth against which the predictions from mixed-cell data were compared.

For each cell-type and locus, the beta-values were grouped into two classes based on sex, and an *F* test of difference of means was performed. The resulting *p*-values were adjusted using the Benjamini-Hochberg procedure (29) to control the false discovery rate. Using the cut-off *α*, a cell-type was labelled as differentially methylated at a given site if the associated adjusted *p*-value was less than *α*. In this case study we used the value *α* = 1 × 10^−4^. While this implies a degree of false positives and false negatives in this set, we considered this to be a reasonable substitute for a perfect ground-truth, especially since the false positive rate should be low given the value of *α*, and that it is unlikely that whole blood methods would detect cell-type differential methylation without it also being detected in cell-sorted data.

#### C.2. Validation

When the actual locations of Cell-type differential methylation for each cell-type are known, it is possible to perform a Receiver-Operator Curve (ROC) analysis for each cell-type, as well as for general differential methylation prediction. The relationship between panel size and False Discovery Rate (FDR) was explored and compared between methods, and panel sizes were chosen so the FDR was 10% *±* 1%.

## Results

### D. Simulation Study

The marginal densities and tabulated summary statistics for the proportion estimates on which the simulated proportion values are based are given in Suppl. Table 1.

As shown in Figure 2 The Orthogonal Effect-Linear Regression (OE-LR) predicted Differential Methylation for all simulation runs of Scenarios 1 and 3, but only 76% and 44% of runs in Scenarios 2 and 4 respectively. OE-LR exhibited poor performance in Scenario 4, indicating that when differential methylation only occurs in a single uncommon cell-type, this is difficult to detect as an orthogonal effect, since the apparent orthogonal effect of a truly cell-type specific effect is proportional to the average cell-type fraction.

**Fig. 2.**
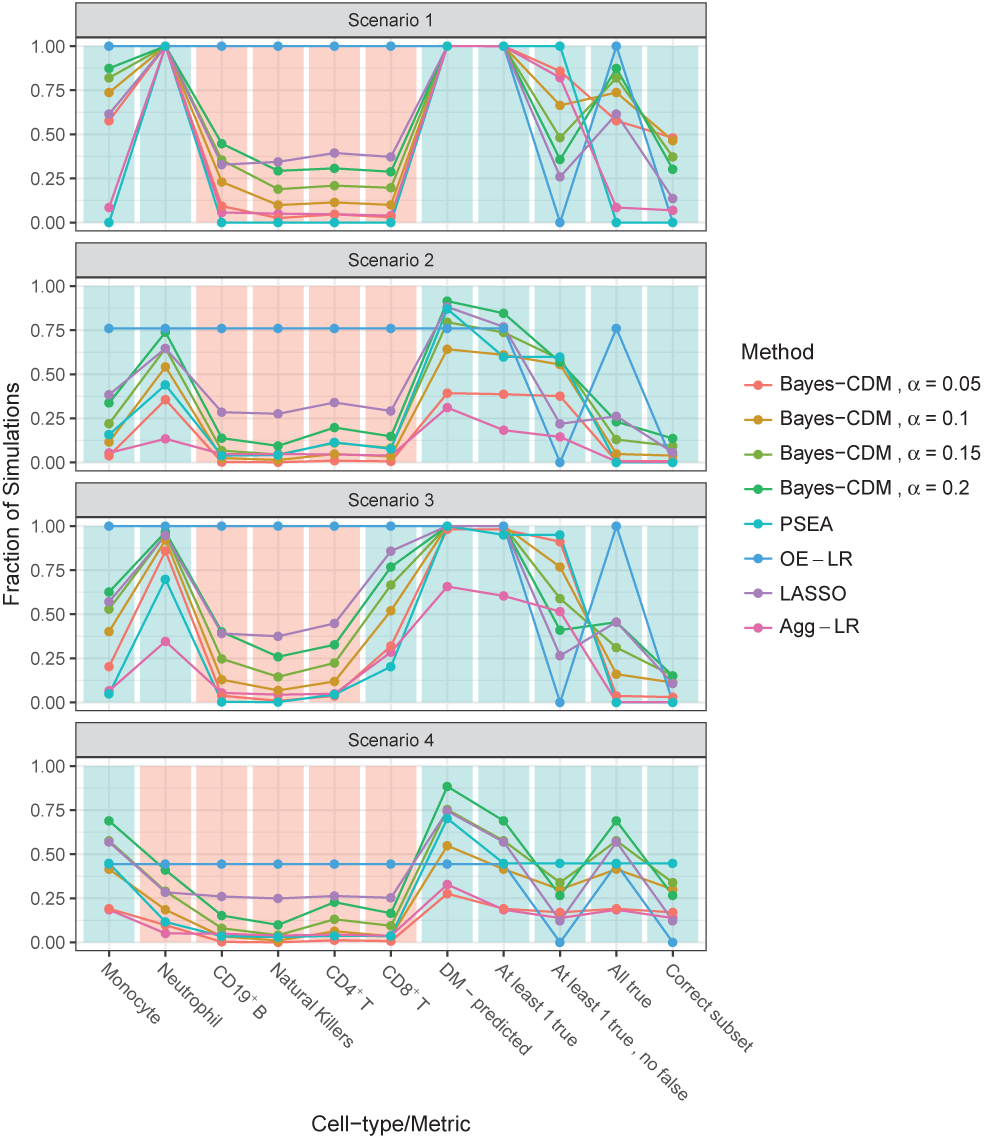
Results for four scenarios with 100 simulation runs each. Bayes-CDM, PSEA, Agg-LR and LASSO methods were used to detect cell-type differential methylation in each of the 6 cell-types, as well as the four metrics. For BayesCDM, several different cut-off values are given. Numbers indicate the proportion of simulation runs where the Cell-type differential methylation was detected or multicell-type metric was met. The rows are coloured blue or red, depending on whether a large or small value should be seen, given which cell-types are differentially methylated in each scenario.

The LASSO method predicted differential methylation at a higher frequency than the other methods in all four scenarios, except for Bayes-CDM at a cut-off value of 0.2. However, the LASSO displayed a tendency to incorrectly predict differential methylation for cell-types where no differential methylation was present. This resulted in a low rate of correct subset prediction. The LASSO performed comparatively well for predicting differential methylation in Scenario 4, indicating that it may be suitable where only a single cell-type is differentially methylated.

The Aggregating Linear Regression (Agg-LR) performed the most poorly in terms of predicting differential methylation and correct subset prediction (excluding OE-LR which is not designed to predict the correct subset). While Agg-LR rarely incorrectly predicted differential methylation, it lacked the sensitivity to predict differential methylation in cell-types where it was present. As a result, Agg-LR predicted the correct subset in 18% of simulation runs in Scenario 3 and ≤8% for the other scenarios. The Agg-LR method was computationally efficient, so it may be useful as a first-pass analysis when the signal is particularly large and from multiple celltypes as in Scenario 1.

The PSEA method tended only to predict differential methylation in a single cell-type if any, which reduced its performance in the scenarios where more than one cell-type was present (1,2, and 4). The method had a very low rate of predicting cell-type level differential methylation incorrectly.

The best method for correct subset prediction was Bayes-CDM, however the optimal cut-off value differed between scenarios. In Scenario 1, a cut-off value of 0.05 was optimal, but in Scenario 4 the cut-off value 0.1 was optimal, and 0.15 was optimal for Scenario 2 and 3. While these values did not correlate with the optimal values for prediction of differential methylation in each scenario, they tracked well with the correct subset measure. This indicates that the prediction of differential methylation increases with cut-off both because of an increase in prediction of truly differentially methylated cell-types, and a higher rate of incorrectly predicting differential methylation. As a result, the rate of correct subset prediction increases as a result of correct prediction, but decreases as the rate of incorrect prediction increases.

For a single simulation run on each scenario, the model in Bayes-CDM was fitted using Hamiltonian Monte Carlo (HMC), and the posterior densities compared with the Laplace approximations. While the true densities displayed some skewness where the Laplace approximations did not, the mean values were close and the overall shapes overlapped well. Therefore we concluded that the Laplace approximations were a sufficiently accurate representation of the true posterior densities (see Suppl. Figure 2).

### E. Case Study: Sex Associations

Based on the cell-sorted data, it was determined that 5813 (1.3%) of CpG loci exhibited differential methylation in at least one cell-type. Of these loci, 2553 (43.9%) exhibited differential methylation in all six cell-types, and were found to be almost all located on the X and Y chromosomes (2538 and 14 respectively; one exception on Chr 3). Of the DMLs with less than 6 differentially methylated cell-types, most of these were also located on the X and Y, although 27 (0.8%) were located on other chromosomes.

A ROC analysis (Figure 3) showed that all methods except for the Agg-LR showed very high discrimination (AUC > 99%) between CpGs with and without differential methylation.

**Fig. 3.**
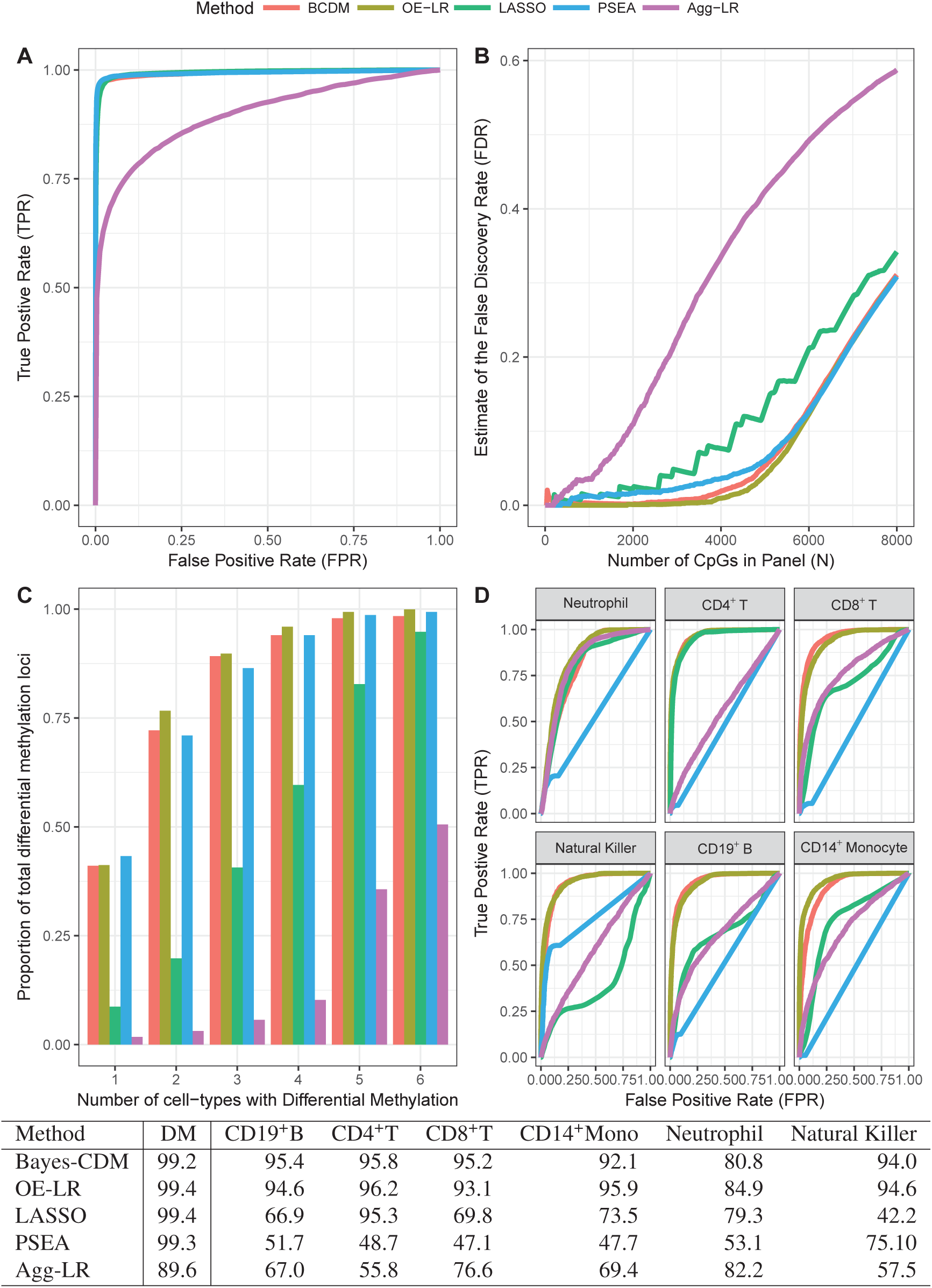
(A) ROC curves for finding Differentially Methylated Loci, showing True Positive Rate (TPR) on the *y*-axis and False Positive Rate on the *x*-axis. Area Under the Curve (AUC) metric given for each method in the legend. (B) False Discovery Rate for panels of predicted DMLs, as it changes with increasing panel size for all 5 methods, calculated using the ground-truth from the cell-sorted data. (C) Frequencies of differentially methylated CpGs found in 10% FDR panels from the five different methods, stratified by the number of cell-types differentially methylated. (D) ROC analysis of the five different methods’ ability to detect differential methylation for each of the six cell-types. AUC (%) values are given in the table below the graph.

**Fig. 4.**
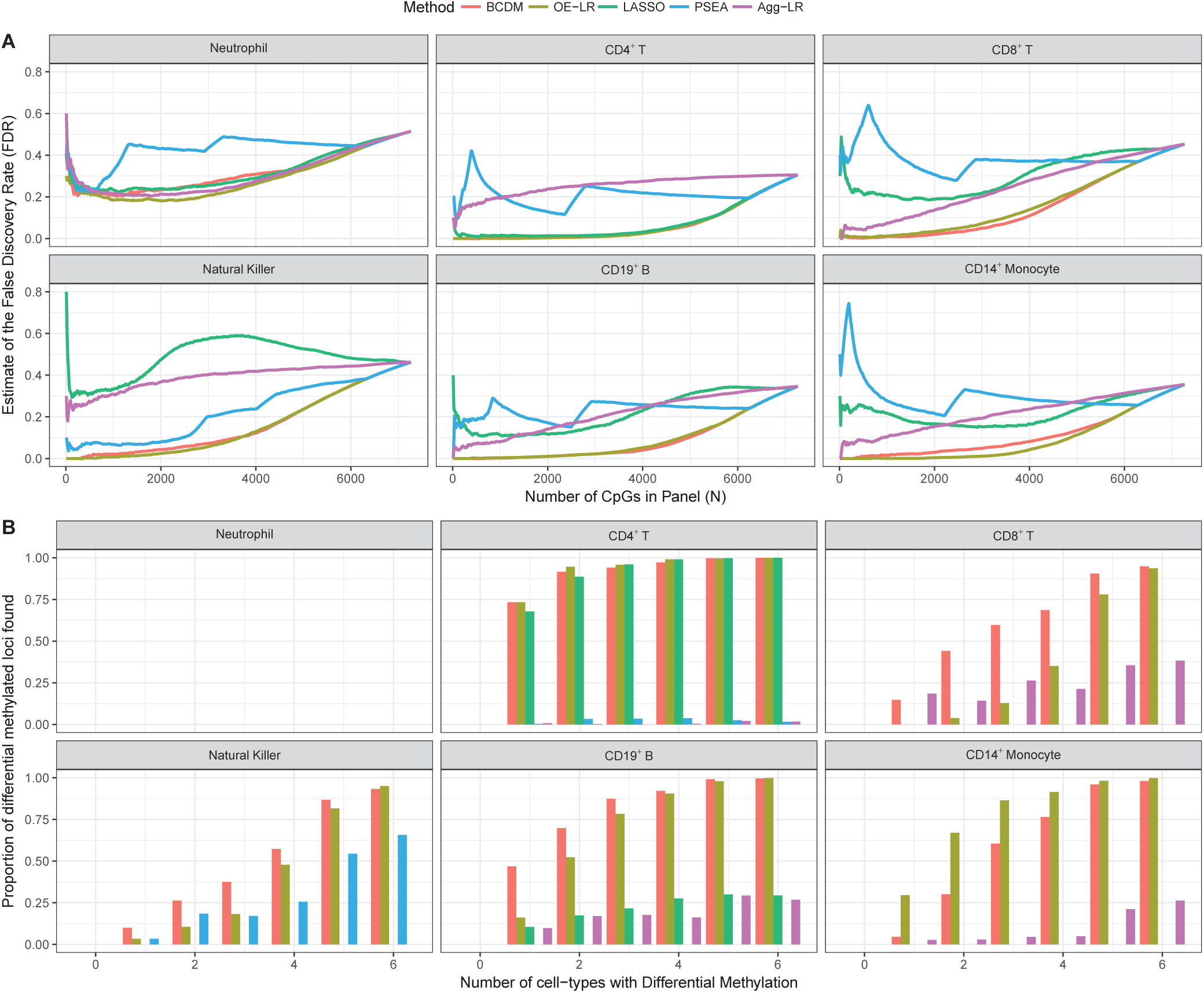
(A) False Discovery Rate (FDR) curve for each method as a function of panel size. Each method creates a different panel for each cell-type. (B) Distribution of CpGs in 10% FDR panels as a function of the number of cell-types with differential methylation. Values here are a proportion of the total number of CpGs with differential methylation for the given cell-type.

The FDR (Figure 3 B) for Agg-LR and LASSO methods grew rapidly compared to the other methods as CpGs were added to the panel, while the FDR for Bayes-CDM, OE-LR and PSEA grew very slowly, indicating these methods were better able to discriminate between differentially methylated sites and non-differentially methylated loci.

Investigating these panels showed that almost all loci which exhibited differential methylation in four or more cell-types were contained within the panels, with the exception of the Agg-LR panel. All panels tended to include fewer loci where less than four cell-types were differential methylated. For all methods, there was indication of a relationship between the number of cell-types with differential methylation at a locus, and the chance of predicting differential methylation at this locus.

### F. Cell-type-specific Differential Methylation Prediction

The second task was to predict differential methylation at loci for specific cell-types. The ROC analysis showed that the Bayes-CDM performed quite consistently over all cell-types. It was the top-performing method for the CD8^+^T and CD19^+^B cell-types, while the OE-LR method performed the best for the Monocyte and Neutrophil cell-types, which tended to be differentially methylated together, resulting in a large effect. For the Bayes-CDM and OE-LR methods, the Neutrophil cell-type appeared to be the most difficult to detect based on the AUC values, despite being the most common cell-type. The PSEA method showed very poor performance for determining cell-type differential methylation for any of the cell-types. The LASSO was generally a poor to moderate performer for the all cell-types except CD4^+^T. The Agg-LR method showed poor comparative performance except for Neutrophil, where it outperformed the Bayes-CDM method. For predicting differential methylation at specific cell-types, the Bayes-CDM and OE-LR performances were almost equal best, although the AUC indicated OE-LR was a slightly better performer.

For panels of the same size the Bayes-CDM and OE-LR methods generally had lower FDR values, and the relationship between FDR and panel size was monotonic, with the exception of Neutrophils where the minimum was at a panel size around 2000. The FDR curve for the PSEA method was not monotonic or smooth, and did not drop below 10% except for Natural Killers.

Panels were selected for a 10 1±% FDR where possible. This produced panels of varying size, including several cases where the FDR was never close enough. Notably no method could produce a panel of any size for Neutrophil with a FDR near 10%. While panels from both Bayes-CDM and OE-LR contained most loci with 5 or more other cell-types also exhibiting differential methylation, Bayes-CDM tended to detect more of the CpG loci with smaller subsets of cell-types exhibiting DM for CD8T, NK, and CD19B, while OE-LR was able to detect more of these loci in Monocytes.

## Discussion

In this paper, we present a new method for predicting differential methylation in specific cell-types when only heterogeneous or mixed cell data are available. The method is based on a Bayesian model which takes account of the inherent constraints on both methylation data and cell-type methylation levels through logit link functions, whilst preserving the linear mixing process assumption. We also incorporate prior knowledge of cell-type relatedness by specifying independent Normal priors on a set of contrast parameters, which act to shrink related cell-type methylation levels together. The method performed relatively well at detecting differentially methylated loci in comparison to other methods, and demonstrated consistent performance in identifying differential methylation associated with phenotype for all cell-types. Almost all methods for predicting differential methylation at the cell-type level require *a priori* cell-type proportion information. The PSEA method (25) uses the mean values of *a priori* known cell-type specific expression as a surrogate for proportion estimates, while the method by Montaño et al. uses reference-based proportion estimate (27) as covariate inputs. An exception is the recent unsupervised method MeDeCom by Lutsik et al. (19), which applies non-negative matrix factorisation in addition to a boundary-weighted regularisation penalty to estimate so-called latent methylation components bounded between 0 and 1, and proportion estimates for each of these components. While this method presents a useful procedure for estimating cell-type methylation level, it does not include phenotype covariate information, and so it is not make clear how to use it for identifying phenotype related differential methylation.

A significant hurdle in modelling the heterogeneous data is that both the beta-value data and the cell-type methylation levels are restricted to the unit interval. A general remedy to the bounded methylation data is to apply a logit-transform, the result being referred to as an *M*-value. The *M*-value is unrestricted, thus allowing standard analysis tools with Normal assumptions to be used more effectively (30). However, a critical issue in the case of mixed cell data is that if methylation is characterized in terms of *M*-values, then the linear mixing process assumption used in previous methods needs to be redefined.

Methods which consider cell-type heterogeneity can model phenotype effects either orthogonal to proportion estimates such as RefFreeEWAS (21), or as interactions between phenotype and proportion, such as PSEA Kuhn et al. (25) and the method by Montaño et al. (18). Kuhn et al. (25) and Montaño et al. (18) give similar mathematical arguments for this conclusion.

Previous methods have used a Normal distribution for beta-values without any constraints (18) or used constrained optimisation (19). (30) showed empirically that standard statistical tests for differential methylation with normal distribution assumptions were more powerful when the input data was the *M*-value version rather than beta-value. This method uses a logit-Normal model, which is equivalent to using a normal distribution with *M*-value data, but the linear mixing process assumption is still valid. Additionally, with the exception of White et al. (13), this is the first method to incorporate prior information from the cell-type lineage to improve inference. Whereas White et al. (13) used the cell-type lineage in a model selection context to identify a small set of candidate models based on lineage groupings, here we use a single model and incorporate the lineage as informative prior distributions.

The shrinkage parameter *λ*_0_ controls the informativeness of the prior, and is analogous to the tuning parameter in ridge regression or other penalized regression methods. Because a hierarchical approach was used where *λ*_0_ is conditionally separated from the parameters of interest by *λ*_1_ and *λ*_2_, this should mean the inferences are somewhat sensitive to choice of *λ*_0_. Nevertheless, the value could potentially be optimised by cross-validation or via information criteria such as AIC or BIC. However, these require multiple runs of the model for different tuning values, which has a high computational burden. An alternative could be the Empirical Bayesian Gibbs Sampler proposed by Casella (31), which uses an EM algorithm step before the Monte Carlo sampling algorithm to optimise the hyperparameter. A related algorithm proposed by Atchadé (32) uses stochastic approximation to optimise the hyperparameter and draw MC samples in the same run. These algorithms have subsequently been employed in several Bayesian regularised regression methods, including the Bayesian LASSO (33), the Bayesian elastic net (34), and the Bayesian adaptive LASSO (35). These MCMC-based procedures are much slower than the optimisation method used, and so present a significant computational challenge given the large number of loci. Despite this, the problem remains embarrassingly parallel.

From the results of our case study, it is evident that the ability of any method to predict differential methylation at a locus depends on the number of differentially methylated cell-types and the cell-type of interest. Among the compared methods Bayes-CDM was close to the best for finding differentially methylated loci, but investigation of the 10% FDR panels showed that Bayes-CDM was more likely to correctly predict differential methylation at loci with a single differentially methylated cell-type for some cell-types. It is therefore recommended that orthogonal methods such as OE-LR or heterogeneity correction methods (e.g. RefFreeEWAS (21) or ReFACTor (23)) are used alongside Bayes-CDM in finding differentially methylated loci. Differentially methylated loci found using the former could subsequently be investigated with Bayes-CDM to predict cell-type differential methylation.

The simulation scenarios showed that Bayes-CDM was broadly better at accurately detecting cell-type differential methylation where differentially methylated cell-types were contained in a lineage grouping. This indicates that the Bayes-CDM model is more suited to loci where the underlying profile of cell-type methylation levels fits well within lineage groupings. The cell-sorted data indicates that a large portion of the methylome is significantly associated with celltype groupings (13), in comparison with the number associated with specific cell-types. Performance of Bayes-CDM at detecting differential methylation restricted to a single celltype is limited for small sample size, but as the amount of information from data increases, the bias in the posterior toward cell-type groupings would decrease, resulting in more precision for specific cell-types.

This model-based approach has a great deal of potential for similar applications where the assumption of a linear mixing process is appropriate. For instance, gene expression array data from mixed cell samples is constrained to the positive real line and can be modelled as a linear combination of underlying cell-type expression profiles (25). A similar model to Bayes-CDM using a log-transform rather than a logit-transform could be applied to detect cell-type specific expression associated with a phenotype. Count-based data from a mixed cell sample could be modelled in a similar way with a binomial, beta-binomial, Poisson, or Poisson-Gamma data distribution.

In the context of tissue for which there is no known level of hierarchy, one could still apply a logit-function to enforce the constraint on the cell-type methylation levels and the whole blood methylation. Instead of applying informative priors on contrast parameters, the priors could be applied to the celltype levels, with a common mean parameter, or another representations based on known external information.

The future work is to expand the model to detect cell-type level differential methylation in multiple neighbouring loci rather than just on a locus-by-locus basis. This will allow investigation of differential methylation regions as well as single loci.

## Acknowledgements

The authors wish to acknowledge Professor David Balding and Professor Terry Speed for their advice regarding modelling constrained data.

## Funding

This research is supported by an Australian Government Research Training Program (RTP) Scholarship, and the Australian Research Council Australian Centre for Mathematical and Statistical Frontiers (ACEMS).

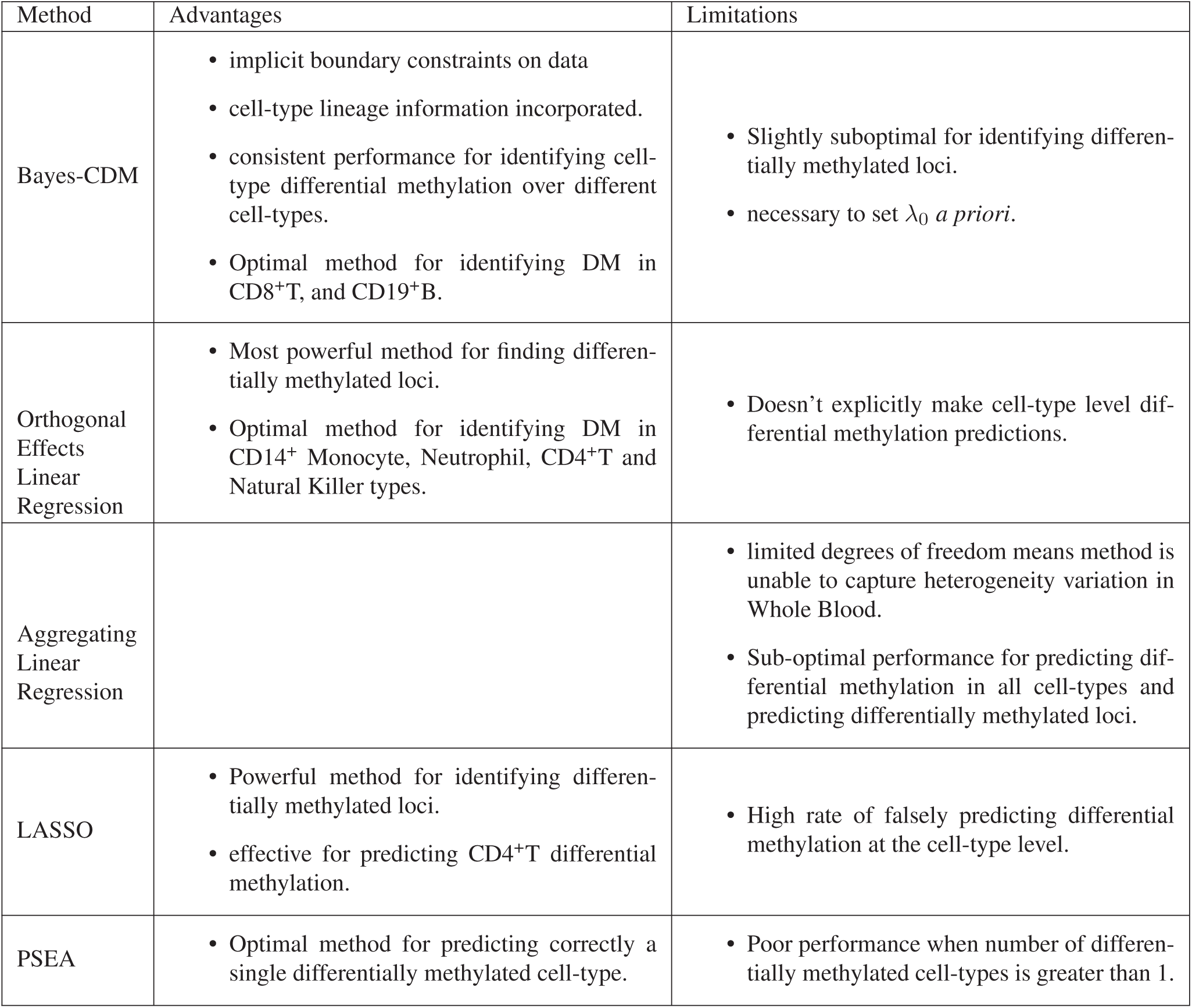

