## Supplementary Notes for "Cell-type specific analysis of heterogeneous methylation signal using a Bayesian model-based approach"

### Supplementary Note 1: Simulation Notes

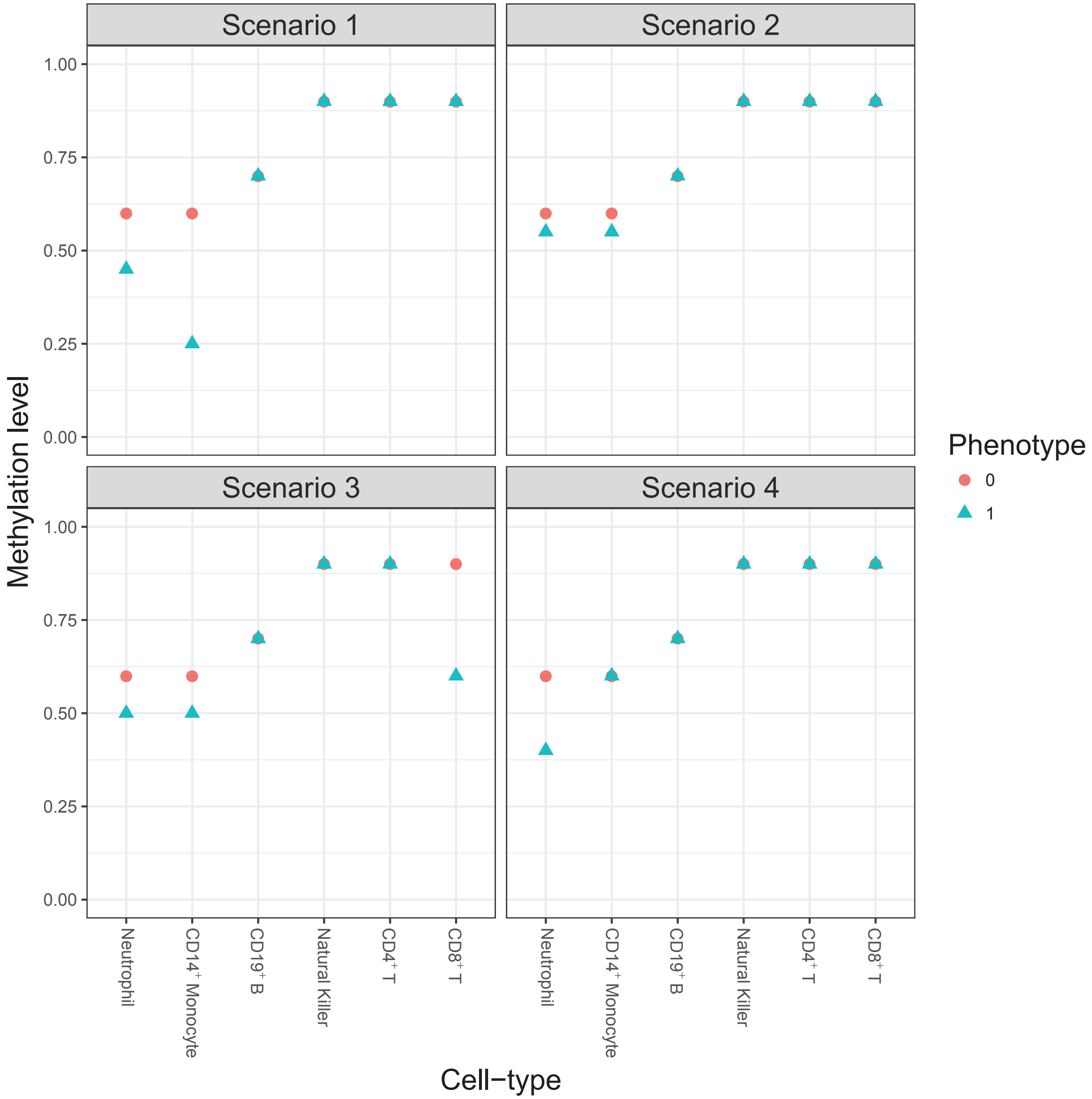


**Supplementary Figure 1.** Simulated cell-type methylation levels (beta-values) depicted for the four simulation scenarios. Shape and colour corresponds to phenotype level.

| | cell_type | mean | sd |
|---|---|---|---|
| 1 | Neu | 0.581 | 0.124 |
| 2 | CD4T | 0.139 | 0.033 |
| 3 | CD8T | 0.076 | 0.035 |
| 4 | NK | 0.027 | 0.011 |
| 5 | CD19B | 0.031 | 0.012 |
| 6 | Mono | 0.081 | 0.062 |

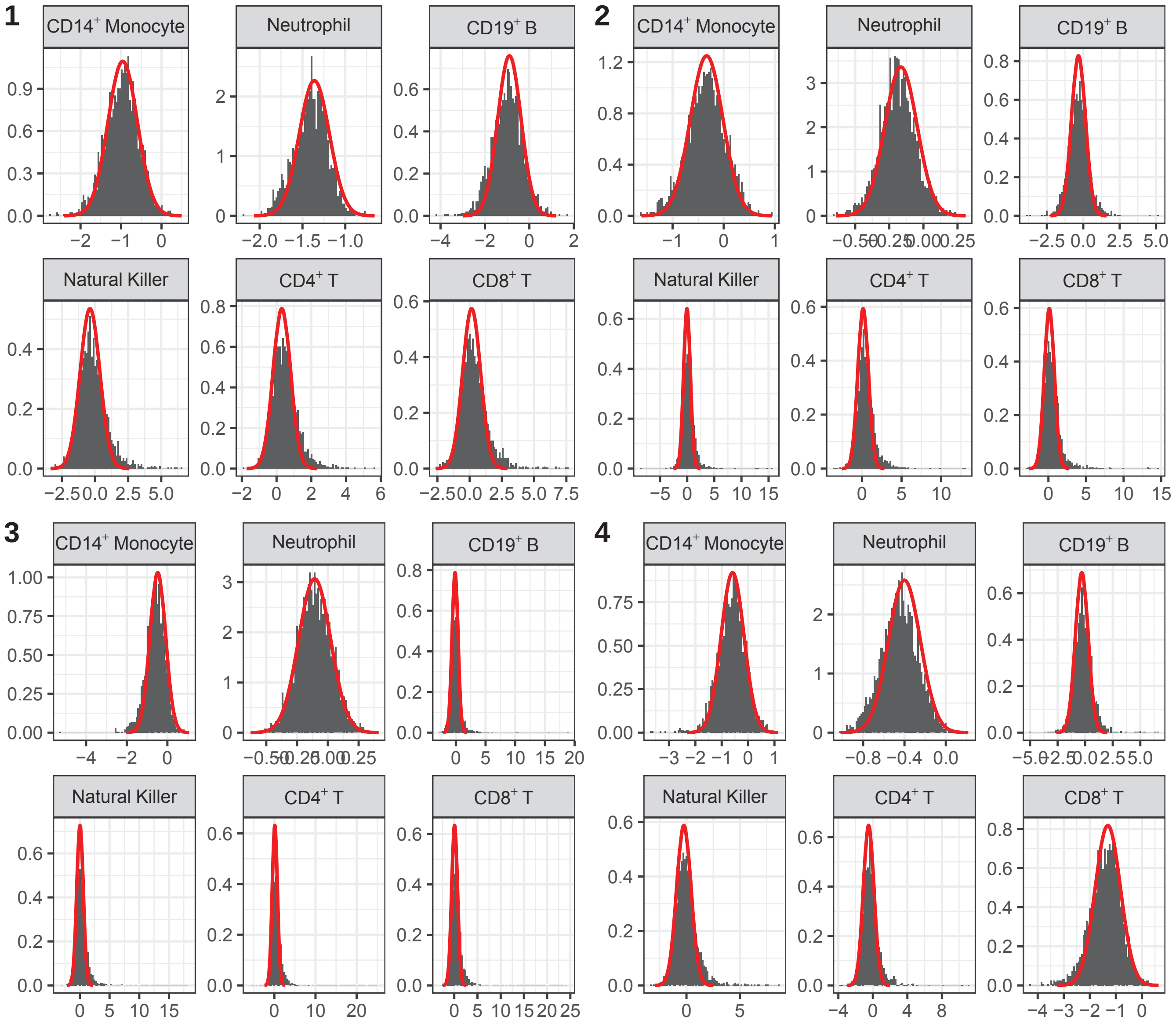


**Supplementary Figure 2.** Comparison between histogram of Hamiltonian Monte Carlo samples and density curve fit derived from the Laplace Approximation (red line)for the $\phi$ parameters.

### Supplementary Note 2: Prior distributions

The lineage matrix and independent priors on $\xi$ parameters were used to create a prior covariance structure representing the cell-type lineage. To ensure this was successful, this was investigated empirically by sampling $\xi$ parameter values directly from the prior distributions and transforming them into $\theta$ parameters. The correlation matrix was then calculated between the $\theta$ parameters, and hierarchical clustering used to cluster them. As expected, the resulting tree resembled the lineage exactly, confirming that the prior information contains the desired covariance structure (see Suppl. Figure 3).

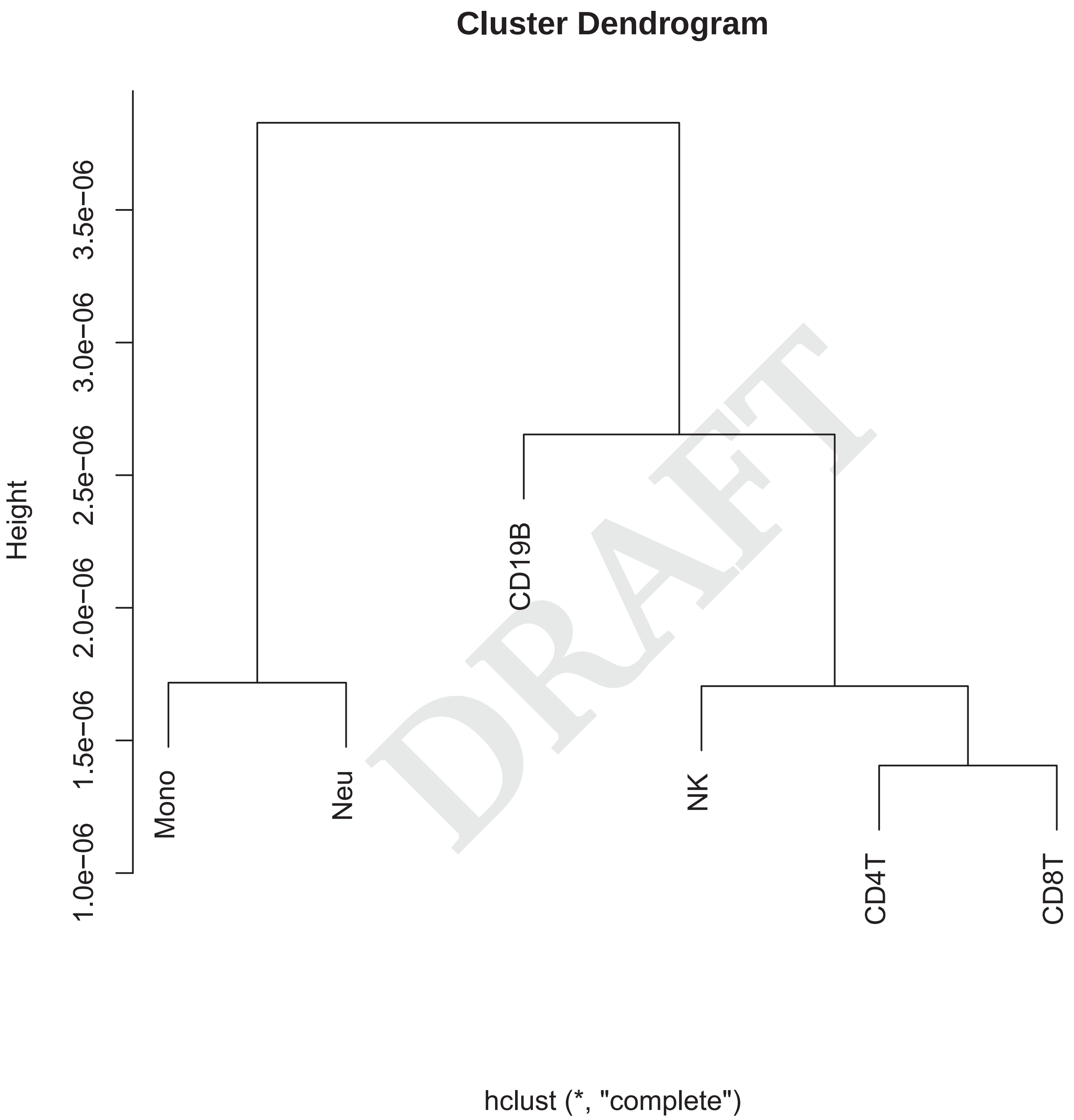


DRAFT

**Supplementary Figure 3.** Heirarchical clustering tree, based on the correlation structure of the samples, which exactly matches the known haematopoietic tree from which the tree matrix is based.

| cell_type | mean | sd | LQ | median | UQ | Q2.5 | Q97.5 |
|---|---|---|---|---|---|---|---|
| Mono | -5.78 | 2293.67 | -10.19 | 0.05 | 10.18 | -125.25 | 122.86 |
| Neu | -5.79 | 2293.68 | -10.17 | 0.04 | 10.19 | -125.37 | 122.83 |
| CD19B | -5.79 | 2293.68 | -10.18 | 0.06 | 10.22 | -125.24 | 122.99 |
| NK | -5.79 | 2293.67 | -10.29 | 0.07 | 10.31 | -125.19 | 122.92 |
| CD4T | -5.79 | 2293.68 | -10.33 | 0.06 | 10.37 | -125.79 | 122.94 |
| CD8T | -5.79 | 2293.67 | -10.35 | 0.06 | 10.37 | -125.34 | 122.93 |

**Table 4.** Summary statistics for the $\theta$ samples drawn from the prior distributions. The statistics did not vary substantially between cell-types, and the large standard deviation, 2.5% and 97.5% quantiles indicate highly spread, and thus weakly informative distributions.

DRAFT
